# Third Harmonic Generation Microscopy Reveals Structure and Mucus Dynamics in Human Airway Epithelium Models

**DOI:** 10.64898/2026.04.10.717621

**Authors:** Doyoung Kim, Alexandra Latshaw, Marta Balkota, Moritz Wiggert, Milvia Alata, Song Hwang, Samuel Constant, Pierre Maechler, Pieter Vanden Berghe, Luigi Bonacina

## Abstract

Airway epithelium plays a major role as the primary interface between human body and the external environment, acting both as a physical and functional barrier. *In vitro* airway models that reproduce the epithelium architecture are therefore a valuable tool for studying infection, inflammation, and transport processes. In this work, we present a label-free, non-invasive method to visualize and measure mucociliary transport in air–liquid human models using third-harmonic generation (THG) microscopy with an optical parametric amplifier laser source at 1300 nm. By exploiting the intrinsic nonlinear contrast at optical heterogeneities, THG provides high-resolution images of both epithelial structures and of the overlying mucus layer without the need for fluorescence staining or sample processing. Time-lapse THG imaging reveals depth-dependent transport dynamics within the mucus, offering new insights into mucociliary transport mechanism. Our approach offers a physiologically relevant way to assess mucociliary function *in vitro* and could support studies on respiratory diseases, drug delivery and efficacy, and epithelial remodeling.

For Table of Contents Only

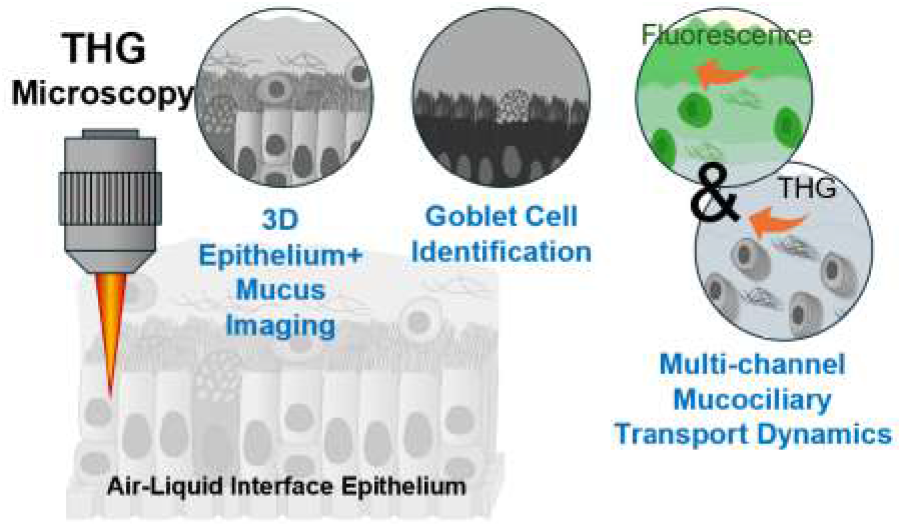

## Introduction

Human-based *in vitro* tissue models have become central to translational research because they can reproduce essential aspects of human physiology better than two-dimensional cultures or animal models, and they are aligned with regulatory trends favoring human-relevant testing approaches. [1] When derived from patient-specific cells, these platforms enable assessment of individual response profiles, prospectively supporting personalized therapeutic strategies. [2, 3]

Within the respiratory system, the airway epithelium plays a major role as the primary interface with the external environment, acting both as a physical and functional barrier. *In vitro* airway models that reproduce the epithelium architecture are therefore a valuable tool for studying infection, inflammation, and transport processes. Air–liquid interface (ALI) cultures stand out among *in vitro* systems because they differentiate into an epithelium that recapitulates key structural and functional properties and are increasingly used for the evaluation of inhaled therapeutics, for pollutant toxicity testing.[4] Finally, their compatibility with patient-derived material make them attractive for translational studies and personalized approaches.

Mucociliary transport (MCT) is the fundamental self-clearing mechanism of the respiratory system, playing a crucial role in maintaining pulmonary health by removing inhaled pathogens and particulate matter from the airways.[5] Impaired MCT is a hallmark of several chronic respiratory diseases, including cystic fibrosis (CF) and chronic obstructive pulmonary disease, and acts as an indicator of disease progression and therapeutic efficacy.[6] Despite its central role in respiratory health and disease, the ability to study MCT in physiologically relevant *in vitro* models remains challenging. Existing imaging modalities often fail to combine high spatial resolution, volumetric readout, and temporal sampling compatible with ciliary and mucus dynamics, while preserving native tissue conditions.[7] Particle-tracking approaches using fluorescent microspheres and widefield fluorescence microscopy are commonly used to quantify mucus transport,[8, 9] but they require exogenous probes that may locally perturb mucus rheology and provide limited insight into depth-resolved flow within the periciliary layer (PCL) and inner mucus regions.[10] Optical coherence tomography (OCT) offers label-free volumetric imaging with micrometer-scale axial resolution and (especially with swept-source implementations) faster acquisition than laser scanning approaches, making it attractive for studies of mucus dynamics.[11–14] As OCT is primarily sensitive to refractive-index discontinuities, it predominantly yields morphological contrast, which can limit its applicability in studies requiring multiplexed optical readouts, such as the integration of fluorescence channels.[15] These considerations motivate the investigation of complementary optical approaches capable of providing simultaneous structural, dynamical, and, when necessary, molecularly sensitive readouts in ALI models, while remaining minimally invasive and compatible with longitudinal studies.

Third harmonic generation (THG) is being increasingly applied as contrast mechanism in microscopy thanks to its high sensitivity to optical heterogeneities and interfaces, and its complementarity with other nonlinear signals such as second harmonic generation (SHG) and multiphoton-excited fluorescence/autofluorescence (MPEF). [16–19] Recent photonics developments have led to the availability of rugged and compact optical parametric amplifier (OPA) laser sources optimized for microscopy, which combine i) short-wave infrared (SWIR) excitation, producing THG signals in the visible range, beyond the cut-off of standard optical components and within the peak sensitivity of silicon detectors, with ii) excitation at low-MHz repetition rates that enhance nonlinear efficiency[20, 21] while minimizing average power and thermal load on samples. Interestingly, recent works have highlighted the possibility of extending THG towards spectroscopic contrast by exploiting sample-specific resonances in combination with multicolor excitation.[22] Owing to its sensitivity to optical discontinuities, [23] THG is particularly well suited to label-free imaging of the air–mucus and mucus–epithelium interfaces. Building on this property, here we employ THG as a label-free, subcellular-resolution modality to visualize the native mucus layer in ALI cultures using a 1 MHz OPA laser tuned to 1300 nm. Because THG resolves individual epithelial cells and reveals fine mucus microstructure (including mucin aggregates), we show that the same contrast mechanism can be used to monitor mucociliary transport dynamics under varying conditions.

## Results and discussion

### Epithelial and Mucus Layer Architecture

As illustrated in Fig. 1a, lung epithelium is primarily composed of a columnar epithelial layer that includes goblet cells (*gob*), basal cells (*bas*), and ciliated cells (*ci*). Each ciliated cell typically possesses 200 to 300 cilia, with each cilium measuring 0.2–0.3 *µ*m in diameter and 6–7 *µ*m in length. Cilia beat at a coordinated frequency, typically of the order of 10-20 Hz, generating a synchronized, wave-like motion that drives directional mucociliary transport across the epithelial surface (Fig. 1b). The epithelium surface is covered with mucus secreted by goblet cells, which traps inhaled particles and pathogens, while ciliary beating facilitates their clearance.[24] The thickness of the mucus layer directly influences ciliary movement and transport efficiency, while abnormal mucus thickness is commonly observed in respiratory pathologies such as CF, asthma, and chronic bronchitis[25].

**Figure 1.**
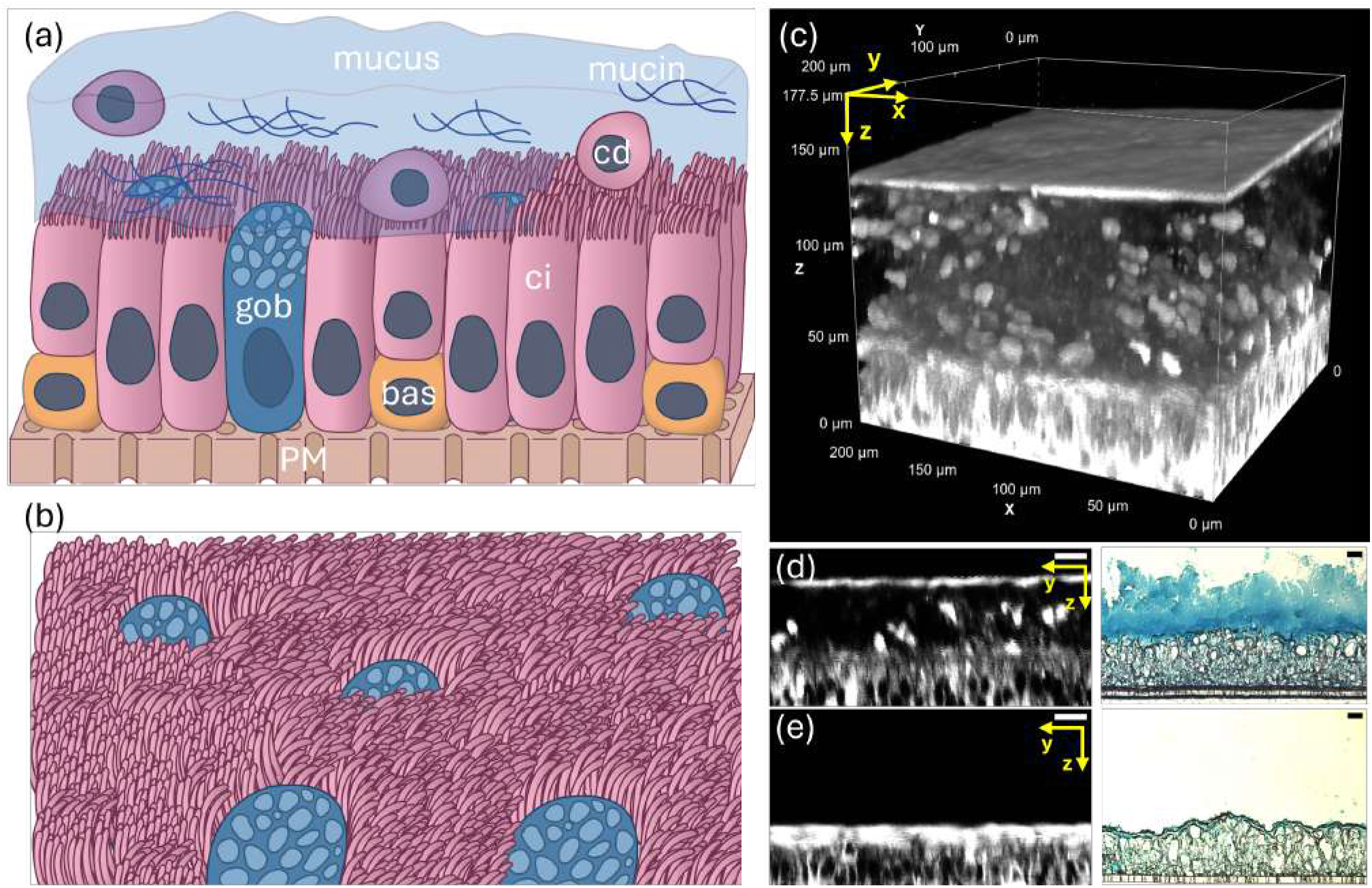
(a) Cross-sectional schematic of the airway epithelium ALI model showing ciliated cells (ci), goblet cells (gob), and basal cells (bas) on a porous membrane (PM). The apical surface is covered by a mucus layer embedding mucins and cellular debris (cd). (b) Illustration of the apical view of the epithelium showing the dense cilia on ciliated cells and the apical secretory region of goblet cells. (c) three-dimensional THG image of an actual ALI sample showing the respiratory epithelial tissue (0-50 *µ*m) and the mucus layer (60-130 *µ*m). (d, e) Cross-sectional THG (left) and histology (right) images of airway epithelium with (d) intact and (e) removed mucus layer. Scale bar: 20 *µ*m.

To investigate the structural organization and functional dynamics of the airway epithelium, we conducted a series of THG imaging experiments on a commercial bronchial ALI model derived from cells collected after surgical polypectomy of informed and consenting patients.

Fig. 1c demonstrates that both the epithelial structure and the overlying mucus layer can be simultaneously visualized by THG. Within the epithelium (*z*: [0 −50]*µ*m), cell nuclei appear as signal-void regions,[26] enabling the identification of individual cells. In the mucus region (*z*: [60 *−*130]*µ*m), THG reveals fine structural details that can be associated with the mucin network and cellular debris (see also Fig. S1). Finally, the mucus upper surface at *z*≃ 130 *µ*m is evidenced by an extremely intense THG signal delineating a rather flat interface with air. The mucus layer thickness, from the epithelial surface to the air–mucus interface, can be estimated in a label-free fashion by THG providing a value in agreement with that obtained on cryosectioned samples stained by Alcian blue[27], as shown in Fig. 1d.

The association of mucus with THG optical contrast is further confirmed by the bulk disappearance of the signal from this region when the mucus layer is removed by rinsing (Fig. 1e). Under these conditions a strong THG signal arises from the newly generated air-epithelium surface, predominantly associated with the sharp refractive-index contrast at the epithelial surface, with a possible contribution from residual mucus remaining after rinsing. To further investigate the impact of the optical properties on cell THG contrast, we filled the apical surface of the epithelium with Phosphate-Buffered Saline (PBS) after fixation. Under these conditions, bundles of cilia on ciliated cells became observable by THG as reported in Fig. S2, which shows the same epithelial region imaged with epithelium exposed to air (Fig. S2a, c) and after apical addition of PBS (Fig. S2b, d). When the epithelium is fixed and immersed in PBS, cilia bundles of 6 *µ*m approximate length become distinct, whereas under direct air exposure, the strong THG signal from the interface with the residual mucus layer dominates, hindering correct assessment of their morphology.

In presence of an unaltered mucus layer, the upper surface of the epithelium appears as in Fig. 2b. A pronounced THG signal marks regions occupied by ciliated cells. By contrast, goblet cells are revealed by the absence of THG emission. In the cross-sectional view (Fig. 2e), goblet cells exhibit a characteristic cup-like morphology in negative contrast, whereas ciliated cells appear bright, except for their nuclei in the basal region of the epithelium. These assignments are corroborated by simultaneously acquired SPY650-Tubulin fluorescence (Fig. 2a, d), which selectively labels cilia and shows strong correspondence with the apical regions of structures associated with ciliated cells.[28] The ability to identify goblet cells through intrinsic nonlinear optical response (also confirmed by a control measurement performed in the absence of staining agents, as in Fig. S3) provides a valuable means to assess their distribution and abundance across different tissues.

**Figure 2.**
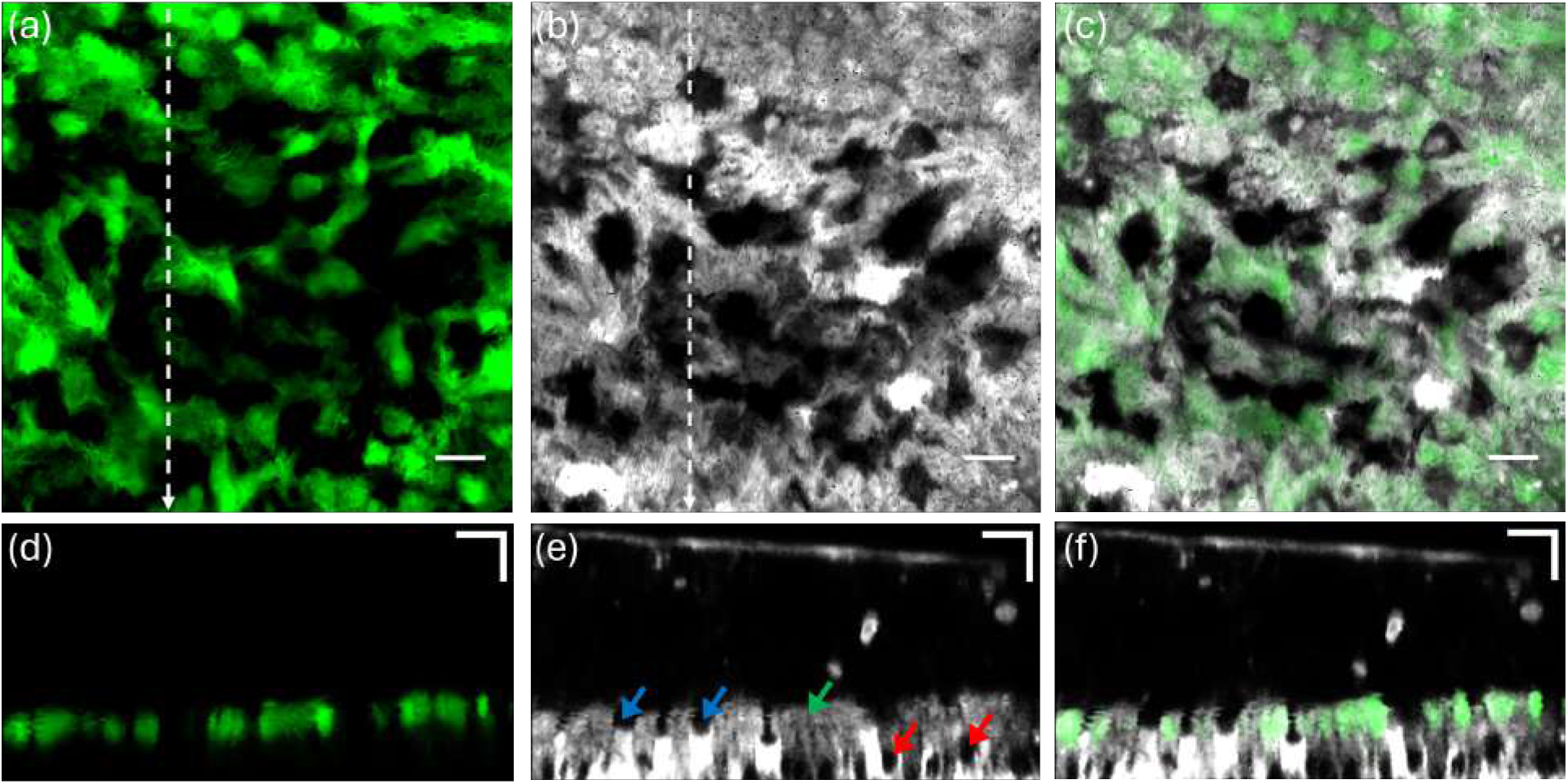
Minimum intensity projection of (a) cilia stained with SPY650-Tubulin, (b) THG signal from the cilia-mucus interface in the same region and (c) merged image. (d–f) Orthogonal views corresponding to the dashed line in (a-b). Goblet cells appear as void within the upper epithelial layer. Nuclei void signals are indicated in red, goblet cell granule void signals in blue, and the cilia–mucus interface signal in green. Scale bar: 10 *µ*m.

### THG-enabled Monitoring of Mucociliary Transport

The presence of multiple THG emitting structures within the mucus layer (such as those marked with red arrows in the orthogonal view in Fig. 3*e*) can be directly exploited for tracing mucus dynamics, as these endogenous objects move in concert with the mucus layer. Panels 3*a-d* show traces of their displacements over a 3.2 min time span (Video 1). The extracted traces indicate that the objects move along the same direction but have a gradient in speed moving from the PCL (panel *a*, 55 *µ*m height) to the upper mucus region (panel *d*, 85 *µ*m height). This height-dependent speed modulation is prominent in the time-resolved orthogonal view shown in Video 2.

**Figure 3.**
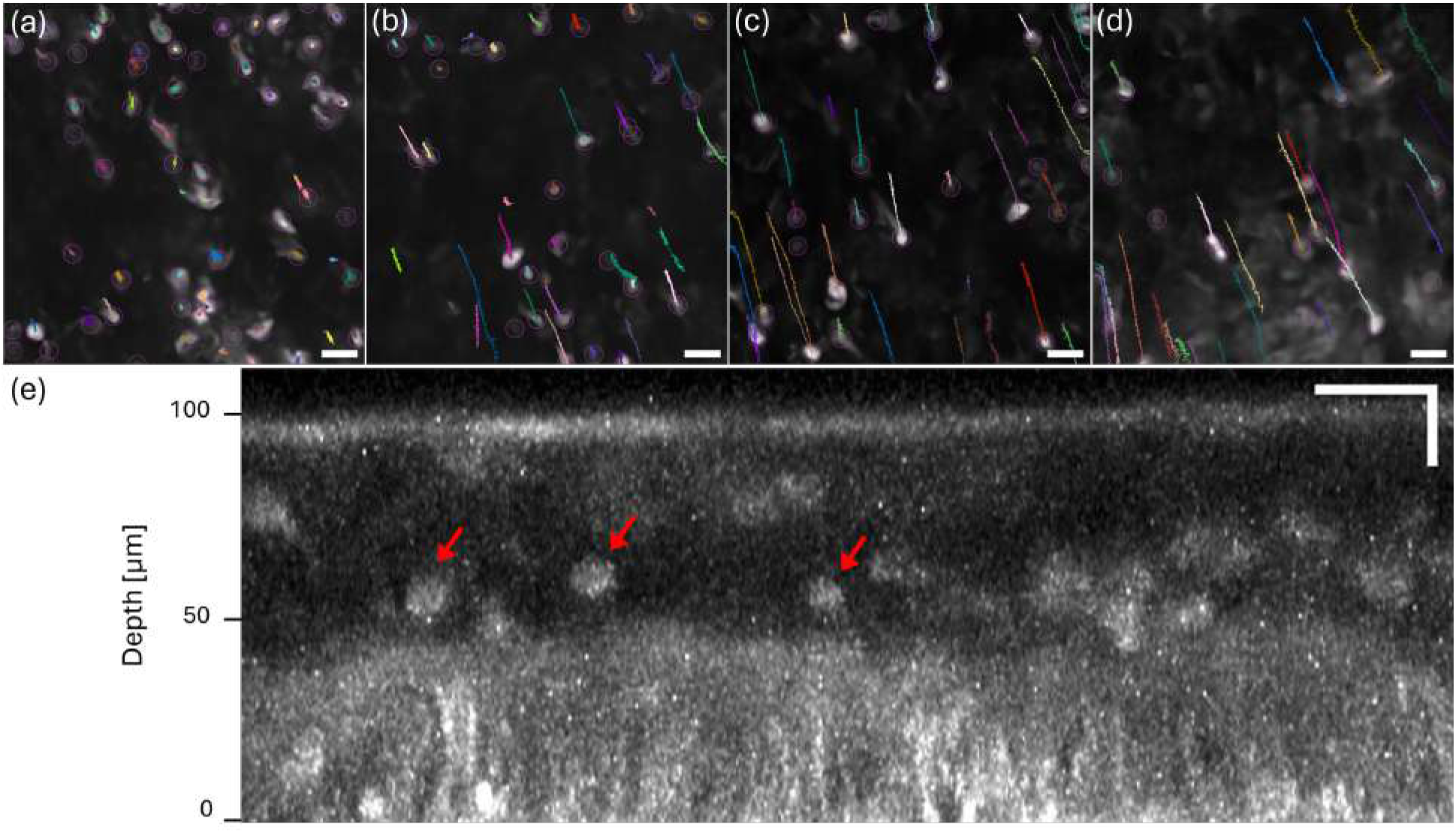
Tracking of cellular debris in the mucus, measured at different heights: (a) 55 *µ*m (near PCL), (b) 65 *µ*m, (c) 75 *µ*m, and (d) 85 *µ*m above the epithelial base. A time-lapse orthogonal view of the same region is shown in Video 2. (e) Its corresponding maximum-intensity orthogonal view of THG signal of mucus and epithelium. Cellular debris within the apical mucus layer are indicated by red arrows. Scale bar: 20 *µ*m.

In the dataset presented in Fig. 3, the observed decrease in lateral speed just above the epithelium can be explained by the mechanical dynamics of ciliary motion: while the ciliary power stroke drives forward transport, the recovery stroke, during which the cilia bend back toward their original position, occurs close to the epithelial surface and locally reduces the effectiveness of forward motion within the PCL.[29] Such a bidirectional interaction introduces localized flow fluctuations that can reduce net transport velocity at the bottom of the layer (Panel a). In contrast, at the interface with air (Panel d), the mucus layer experiences reduced mechanical constraints and is less influenced by the recovery stroke dynamics, which are largely confined to the periciliary region near the epithelial surface, resulting in more uniform bulk transport. [30] Consistent with this interpretation, numerical simulations resolving the airway surface liquid as a two-layer system predict markedly reduced velocities within the PCL compared to the overlying mucus layer.[31, 32] Moreover, airway mucus exhibits a pronounced vertical organization, consisting of a low-viscosity PCL adjacent to the epithelium and a denser, more viscoelastic gel-like mucus layer above. In healthy conditions, the upper mucus layer displays greater structural cohesion due to its higher mucin concentration, enabling more efficient transmission of ciliary propulsion and entraining embedded components, such as cellular debris, within the moving mucus bulk. [33, 34].

The mucus transport velocities observed in this study are generally lower than those reported in the literature, which themselves span an exceptionally broad range (up to several mm/s), that can be attributed to the substantial variability in sample characteristics and conditions. In a set of measurements using samples from 3 donors, we measured transport speeds ranging from 1.2 to 2.7 *µ*m*/*s after averaging on multiple individual tracks obtained by following debris over several tens of seconds at a fixed height in the mucus. In some of these tracks we observed speed spikes up to 15 *µ*m*/*s. Although previous studies have reported that ALI models generally exhibit slower transport rates than *in vivo* tissues,[35] and that the motion of endogenous mucus is typically reduced compared to that inferred from tracer bead tracking,[36, 37] the discrepancy observed here remains appreciable. We do not necessarily attribute this difference to sample dehydration or tissue damage, as (i) no decrease in mucus layer thickness was detected over time and the epithelial architecture remained morphologically intact throughout the measurements. The observed velocities are also not constrained by the acquisition speed of our system. Considering a 200× 200 *µ*m^2^ field of view and 1.55 Hz frame rate and assuming four consecutive detections of the same structure for reliable tracking, the maximum measurable velocity is estimated to 70 *µ*m*/*s. Within this work, we did not further pursue absolute quantitative speed investigations in multiple samples, as the primary objective is not to establish reference values for MCT, but rather to demonstrate the imaging capabilities of the proposed THG-based approach applied to human airway epithelial models.

The label-free access to volumetric mucus dynamics enables the investigation of MCT in ALI models affected by specific respiratory conditions and their real-time response to *stimuli*. This approach is demon-strated in Fig. 4 and Supplementary Video 1, where increasing PBS volumes were applied to the apical surface of an epithelium derived from a CF patient. We clearly observe that CF mucus, known for being abnormally thick and viscoelastic, responds strongly to even modest dilutions. In the absence of PBS (Panel a), mucus movement is minimal, consistent with stagnant transport. However, following PBS application, MCT velocity increased dramatically (*∼* 11.6-fold) 6 hours after application of 10 *µ*L of PBS (Panel c). These results highlight the potential of THG-monitoring for pharmacological applications both for assessing topical therapies for respiratory diseases and for studying inhalation-based delivery routes for systemic drug absorption through the airway epithelium.

**Figure 4.**
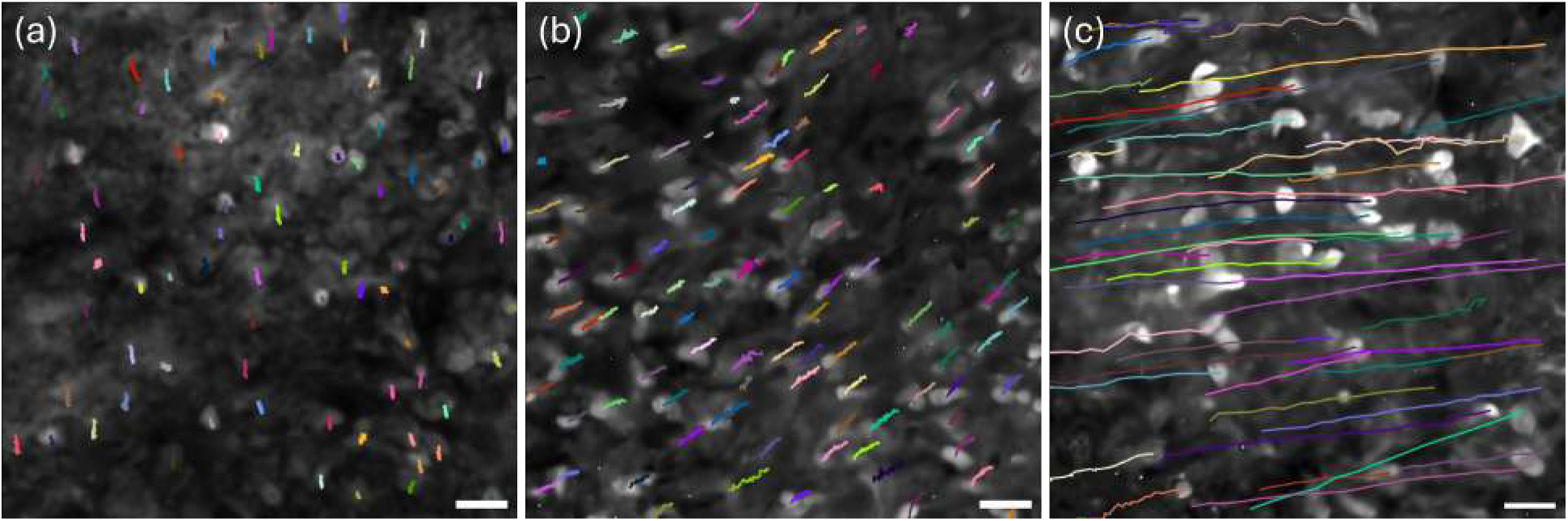
Maximum intensity projection of the mucus layer of epithelial tissue acquired 6 h after PBS application. The epithelium was cultured with primary bronchial cells from a patient with CF, where (a) 0 *µ*L, (b) 5 *µ*L, and (c) 10 *µ*L of PBS were applied apically. Scale bar: 20 *µ*m.

To gain further mechanistic insight into how osmotic agents modulate MCT, we applied a 6 % NaCl solution supplemented with Sulfo-Cyanine5 (Cy5) dye to the apical surface of a CF model with a nebulizer (Fig. S4) and monitored the effect on MCT over time. As reported in Fig. S5 and Supplementary Video 2, the approach allows tracking the diffusion dynamics of hypertonic saline through the mucus layer and monitor the progressive restoration of MCT over time. Initially, mucus and mucins exhibit minimal movement consistent with CF conditions (Panel a). Upon deposition of the solution (Panel b), the hypertonic saline rapidly penetrates and diffuses through the mucus layer within 2 minutes, restoring MCT (Panel c,d). MCT enhancement was observed immediately after diffusion, in agreement with the rapid MCT improvement reported upon hypertonic saline treatment in CF.[38] While previous investigations could show that nebulized hypertonic saline enhances mucociliary clearance in CF,[39] the nonlinear (THG and MPEF) approach demonstrated here allows simultaneous, high-resolution visualization of both NaCl diffusion and mucus movement.

These results gain further significance when compared to widely adopted experimental approaches, where MCT is typically assessed by fluorescent bead tracking under widefield microscopy. While this method provides insights into surface-level mucus dynamics, limitations have been reported.[40] Larger beads (typically >10 *µ*m in diameter) are often employed to ensure visibility and ease of tracking; however, their size restricts them to the mucus surface, preventing penetration into deeper layers and thereby offering limited information about inner-layer dynamics. Smaller beads (1 *µ*m or less) have the potential to probe deeper into the mucus and sample pore-scale microenvironments but they tend to be weakly coupled to large, cohesive mucus assemblies and therefore may move independently of bulk mucus transport.[36, 41] Moreover, because fluorescent bead assays typically involve applying an exogenous bead suspension to the airway surface, the added liquid can locally dilute or transiently rehydrate the mucus layer, altering its concentration and rheological properties and transiently biasing MCT measurements.[42] As a benchmark experiment, 1 *µ*m fluorescent beads (FBs) were applied to the mucus surface and imaged two hours later (Supplementary Video 3). Prior to bead application, the mucus exhibited largely unidirectional transport (Panel a). Following bead deposition, both FBs and endogenous mucus components displayed more disorganized, multidirectional motion (Panel b). This temporary disruption likely arises from the bead application procedure itself, rather than a persistent alteration of mucociliary function. Indeed, a pre–post analysis (Supplementary Video 4) shows that approximately 20 hours after bead application, MCT stabilizes and recovers its unidirectional character. A similar behavior was observed on *N* = 3 samples.

## Conclusions

THG microscopy is a powerful nonlinear imaging modality, highly sensitive to optical interfaces and microscopic heterogeneities.[16, 20] Its application to physiologically sensitive human-derived systems has recently become increasingly viable with the advent of SWIR femtosecond OPAs operating at low-MHz repetition rates.[20, 21, 43] By concentrating excitation energy into fewer pulses, these sources deliver the high peak powers required to drive higher-order nonlinear processes while maintaining modest average power, in contrast to conventional titanium-sapphire and optical parametric oscillator (OPO) lasers operating at 80 MHz. At the same time, OPAs remain compatible with microsecond dwell time scanning regimes, ensuring the possibility of acquiring volumetric imaging of dynamical systems. The advantage of the low-MHz excitation regime is demonstrated in Fig. S6 and Supplementary Video 5, where an ALI model was imaged using both an *∼*80 MHz OPO laser and the 1 MHz OPA system. The OPO required an average power of 36.0 mW to generate sufficient THG contrast for qualitative comparison with the OPA operating at 2.1 mW.[44] As shown, the use of the 80 MHz system led to a significant reduction of the mucus layer thickness over 21 minutes, most likely due to evaporation. Morphological changes in the epithelium and the appearance of damage-associated emission were also observed.

THG imaging applied to ALI *in vitro* models of human airway epithelium provides direct access to structural and dynamical information without the need to process the samples or introduce exogenous agents. This represents a clear advantage, as it preserves the system in conditions close to its natural state and enables longitudinal measurements over time with minimal perturbation. We show that THG contrast allows clear delineation and quantification of the mucus layer thickness covering the epithelium. Within the epithelial layer, characteristic THG morphological features permit discrimination among different cell types. Furthermore, by tracking the position of objects naturally present within the mucus, it is possible to monitor mucus transport as a function of three-dimensional localization, with micrometer spatial resolution.

Notably, THG imaging can be readily combined with complementary optical readouts to extend its range of applications. Its ability to track MCT dynamics under external *stimuli* makes it a valuable tool for drug screening, when combined with fluorescently labeled compounds.[45, 46] Similar multi-channel strategies can be applied to investigate epithelial remodeling (*via* SHG to monitor fibrillar collagen)[47] and mucus abnormalities in disease models such as CF and chronic bronchitis. The potential of THG for short- and long-term monitoring of human 3D airway models is particularly compelling given current regulatory shifts toward human-based systems in preclinical and drug development studies, in full ethical alignment with the 3R principles advocating the reduction and refinement of animal experimentation.[48]

## Experimental

### Culture of Human 3D Lung Epithelial models

Commercially available primary human airway epithelial cultures (MucilAir^™^, Epithelix Sàrl, Geneva, Switzerland) were used in this study. These cells were obtained from patients undergoing surgical polypectomy. All experimental procedures were fully explained to participants, and informed consent was obtained from all donors. The study was conducted in accordance with the Declaration of Helsinki (Hong Kong amendment, 1989) and received approval from the *Commission Cantonale d’Éthique de la Recherche Scientifique de Genève* (reference number 15-062). All samples were collected with informed consent as part of ethically reviewed and approved protocols. Collection procedures included (i) obtaining detailed donor medical histories to ensure the safety of both donors and researchers, and (ii) approval by an Institutional Review Board (IRB). Primary human bronchial and nasal epithelial cells were isolated from biopsies provided by consenting donors. The epithelial cells were cultured on the microporous membrane of a Transwell insert (Oxyphen, Wetzikon), and after several days, were transitioned to air-liquid interface (ALI) conditions for at least 28 days to allow full differentiation into a ciliated epithelium. Basolateral medium (EP04MM, Epithelix Sàrl; 700 *µ*L) was replenished twice weekly. For co-culture experiments, lung fibroblasts were seeded onto the basolateral side of a 6.5 mm Transwell insert containing differentiated epithelial cells and allowed to attach for 3 hours. Throughout the culture period, inserts were maintained at 37 °C in a 5 % CO_2_ humidified atmosphere and handled strictly under sterile conditions. During image acquisition, the tissue was maintained at 37 °C in a mini incubator (H501-T, Okolab), con-nected to a humidifying pump to ensure a stable environment. Each imaging session lasted no longer than 5 hours, after which the tissue was discarded to prevent any risk of contamination. The donor information for each experiment is given in Table 1 in Supporting Information. Donor A was a 61-year-old female, African American and non-smoker, donor B was a 22-year-old male, Caucasian, non-smoker. Donor C was a 22 years old male, Caucasian, non-smoker and donor D was a 59-year-old male, Caucasian, non-smoker and donor E was a 27-year-old male, Caucasian, non-smoker. Donor F was 35-year-old male, origin not reported, non-smoker and donor G was 59-year-old female, Caucasian, non-smoker. Donor H was a a 22-year-old male, Caucasian, non-smoker. Donor I was a 72-year-old male, Hispanic, non-smoker. Donor J was a 72-year-old male, Hispanic, non-smoker. Details for donor K were not reported. Donor L was a 59-year-old female, Caucasian, non-smoker. Donor M was a 30-year-old female, origin not reported, non-smoker and had Homozygote ΔF508 mutation which is the most common mutation causing CF. Donor N was a 39-year-old male, origin not reported, non-smoker and had ΔF508 mutation.

### Sample preparation

**Figure 1d and e**. THG images are compared with histology sections of adjacent tissue regions. For mucus removal, 20 *µ*L of 0.9% NaCl solution was added to the apical side of the inserts, followed by incubation at 37 °C for 20 minutes. The solution was then entirely removed after thorough mixing with a pipette. Subsequently, the inserts were fixed by complete immersion in 4% paraformaldehyde (PFA) in PBS for 20 minutes at room temperature. The remaining inserts were fixed by immersing the basolateral side in 1 mL of 4% PFA for 20 minutes, without adding any medium to the apical side to preserve the native mucus layer. After fixation, tissues were embedded in OCT compound (Cellpath, KMA-0100-00A) and immediately frozen at –20 °C. Sections were then cut at 10 *µ*m thickness using a cryostat (Leica CM3050 S) and mounted onto glass microscope slides. The microscope slides were subsequently stained with Alcian blue (Sigma-Aldrich, B8438) for 10 minutes, then thoroughly washed and mounted.

**Figure 2a and d**. SPY650-Tubulin (Spirochrome SC503) was diluted 1:1000 in media and added to the basolateral side of MucilAir, followed by a 48-hour incubation before imaging.

Mucus transport speed analysis was conducted on samples derived from donors A, B, and D.

**Supplementary Video 2** Sulfo-Cyanine5 carboxylic acid (13390, Lumiprobe) was diluted in 6 % NaCl (Mucoclear 6 %, PARI) to a final concentration of 1 *µ*g/ml, then nebulized and applied to the apical side of the MucilAir insert using a nebulizer (NE-U100, Omron). The configuration of the nebulization setup is shown in Fig. S4 of the Supporting Information.

**Supplementary Video 3 and 4** Fluorescent beads with a mean particle size of 1 *µ*m (L4655-1ML, Sigma Aldrich) were diluted 1:500 in PBS before being applied to the MucilAir insert. A volume of 3 *µ*l of the diluted solution was deposited onto the insert using a pipette, then the insert was kept in the incubator at 37 °C and 5 % CO_2_. Donors J and K were additionally included in the fluorescent bead assay for 2 h monitoring, and donors K and L were additionally included for the 24 h recovery monitoring.

### Nonlinear Microscopy and Image Analysis

The THG signal from epithelial tissue was generated at 1300 nm using the Short-Wave Infrared Microscope (SWIM) at the University of Geneva. SWIM combines a commercial microscope matrix with custom SWIR-optimized optics (Miltenyi Biotec, Bielefeld, Germany) and a CRONUS-3P laser (1 MHz OPA, 1250–1800 nm; Light Conversion, Vilnius, Lithuania). The experimental configuration and sample placement are illustrated in Fig. S7. The system was equipped with an Olympus XLPLN25XWMP2 water-immersion objective (25×, NA 1.05, WD 2 mm). THG emission was detected using a GaAsP-photocathode PMT (Hamamatsu H12056-40) with a 450/60 nm bandpass filter. Fluorescent signal (SPY650-tubulin, Cy5 emission) was collected with an infrared-extended multialkali-photocathode PMT (Hamamatsu H12056-20) and a 650/50 nm bandpass filter. The nominal pulse duration of the CRONUS-3P at 1300 nm is 50 fs at the laser output. Group delay dispersion (GDD) pre-compensation was optimized by maximizing the SHG and THG signals from inorganic nanoparticles[49] dispersed on a microscope slide, yielding an optimal setting of −3.6 kfs^2^. A beam shaper was used to expand the CRONUS-3P beam to overfill the back aperture of the objective to maximize effective NA and resolution.

For the comparison shown in Fig. S6, an Insight X3 laser set to 1300 nm (80 MHz OPO; *∼*130 fs pulse duration at laser output at 1300 nm, assuming transform-limited Gaussian pulses; Newport/Spectra-Physics, Milpitas, CA, USA) was aligned through the same microscope platform. No additional beam shaping or external dispersion optimization was performed for the InSight source during these measurements. The average power was measured after the microscope objective (without immersion liquid) using an Ophir 3A-PF-12 thermal sensor and StarBright meter.

Three-dimensional images were obtained by Z-stacking a 2D collection of consecutive scans with 1016 ×1016 pixels and a step size of 0.5 *µ*m (Fig. 1, 2, S2). For time-lapse imaging, three-dimensional data was captured with 516 × 516 pixels and a frame time of 0.644 s with a step size of 2 *µ*m (Fig. 3, 4 and Video 1, Supplementary Video 1, 3 and 5), a step size of 3 *µ*m (Supplementary Video 4) and a step size of 4 *µ*m (Supplementary Video 2). Z-height and XY drift were corrected in Supplementary Video 2, as mechanical vibrations and movements were caused by the nebulizer positioned on the stage when operated. Image denoising for Fig. 1, 2 and S1, S2, S3 was performed using Nikon NIS-Elements software. Orthogonal views were obtained by maximum-intensity projection within a defined ROI along one lateral dimension for each time point and z-slice using a custom-written code in Python. Cellular debris were tracked using the Fiji plugin *Track-Mate*.[50] Depending on image quality and object appearance, different tracking parameters were applied. The built-in *Laplacian of Gaussian* (LoG) detector was used with estimated object diameters ranging from 5 to 15 *µ*m and quality thresholds between 5 and 15. In some cases, manual tracking was performed to ensure accuracy (Video 1 and Supplementary Video 1,3,4). The effective scanning speed used for velocity calculations was extracted directly from the microscope acquisition settings.

## Data availability

All data supporting the findings of this study are available within the paper and its Supplementary Information. Additional raw image stacks and analysis outputs are available from the corresponding author upon reasonable request.

## Code availability

Custom analysis scripts used to process orthogonal THG volumes are available from the corresponding author upon request.

## Acknowledgements

We gratefully acknowledge the funding from the European Project FAIR CHARM (H2020 LEIT Information and Communication Technologies 101016457; http://www.faircharm.eu).

## Supporting information

The following files are available free of charge.

- Supporting Information: Additional THG imaging data and control measurements (Figures S1–S7), donor information table, nebulization setup, comparison of laser excitation conditions (PDF).
- Supplementary videos illustrating mucociliary transport dynamics (PDF, MP4, AVI).

## References

(1) U.S. Food and Drug Administration FDA Announces Plan to Phase Out Animal Testing Requirement for Monoclonal Antibodies and Other Drugs, 2024.

(2) Vlachogiannis, G.; Hedayat, S.; Vatsiou, A.; Jamin, Y.; Fernández-Mateos, J.; Khan, K.; Lampis, A.; Eason, K.; Huntingford, I.; Burke, R., et al. Science 2018, 359, 920–926.

(3) Fisher, C. R.; Mba Medie, F.; Luu, R. J.; Gaibler, R. B.; Mulhern, T. J.; Miller, C. R.; Zhang, C. J.; Rubio, L. D.; Marr, E. E.; Vijayakumar, V., et al. Cells 2023, 12, 2639.

(4) BéruBé, K.; Aufderheide, M.; Breheny, D.; Clothier, R.; Combes, R.; Duffin, R.; Forbes, B.; Gaça, M.; Gray, A.; Hall, I., et al. Alternatives to laboratory animals 2009, 37, 89–141.

(5) Wanner, A.; Salathé, M.; O’Riordan, T. G. American journal of respiratory and critical care medicine 1996, 154, 1868–1902.

(6) Knowles, M. R.; Boucher, R. C., et al. The Journal of clinical investigation 2002, 109, 571–577.

(7) Peabody, J. E.; Shei, R.-J.; Bermingham, B. M.; Phillips, S. E.; Turner, B.; Rowe, S. M.; Solomon, G. M. American Journal of Physiology-Lung Cellular and Molecular Physiology 2018, 314, L909–L921.

(8) Howes, A.; Rogerson, C.; Belyaev, N.; Karagyozova, T.; Rapiteanu, R.; Fradique, R.; Pellicciotta, N.; Mayhew, D.; Hurd, C.; Crotta, S., et al. American Journal of Respiratory Cell and Molecular Biology 2024, 71, 282–293.

(9) Sears, P. R.; Bustamante-Marin, X. M.; Gong, H.; Markovetz, M. R.; Superfine, R.; Hill, D. B.; Ostrowski, L. E. Biophysical Journal 2021, 120, 1387–1395.

(10) Atanasova, K. R.; Reznikov, L. R. Respiratory Research 2019, 20, 261.

(11) Oldenburg, A. L.; Chhetri, R. K.; Hill, D. B.; Button, B. Biomed. Opt. Express 2012, 3, 1978–1992.

(12) Liu, L.; Chu, K. K.; Houser, G. H.; Diephuis, B. J.; Li, Y.; Wilsterman, E. J.; Shastry, S.; Dierksen, G.; Birket, S. E.; Mazur, M., et al. PloS one 2013, 8, e54473.

(13) Chu, K. K.; Unglert, C.; Ford, T. N.; Cui, D.; Carruth, R. W.; Singh, K.; Liu, L.; Birket, S. E.; Solomon, G. M.; Rowe, S. M., et al. Biomedical optics express 2016, 7, 2494–2505.

(14) Pieper, M.; Schulz-Hildebrandt, H.; Mall, M. A.; Hüttmann, G.; König, P. American Journal of Physiology-Lung Cellular and Molecular Physiology 2020, 318, L518–L524.

(15) Drexler, W.; Liu, M.; Kumar, A.; Kamali, T.; Unterhuber, A.; Leitgeb, R. A. Journal of biomedical optics 2014, 19, 071412–071412.

(16) Débarre, D.; Supatto, W.; Pena, A.-M.; Fabre, A.; Tordjmann, T.; Combettes, L.; Schanne-Klein, M.-C.; Beaurepaire, E. Nature methods 2006, 3, 47–53.

(17) Tai, S.-P.; Lee, W.-J.; Shieh, D.-B.; Wu, P.-C.; Huang, H.-Y.; Yu, C.-H.; Sun, C.-K. Optics express 2006, 14, 6178–6187.

(18) Weigelin, B.; Bakker, G.-J.; Friedl, P. Journal of Cell Science 2016, 129, 245–255.

(19) Genthial, R.; Beaurepaire, E.; Schanne-Klein, M.-C.; Peyrin, F.; Farlay, D.; Olivier, C.; Bala, Y.; Boivin, G.; Vial, J.-C.; Débarre, D., et al. Scientific reports 2017, 7, 3419.

(20) Bakker, G.-J.; Weischer, S.; Ferrer Ortas, J.; Heidelin, J.; Andresen, V.; Beutler, M.; Beaurepaire, E.; Friedl, P. Elife 2022, 11, e63776.

(21) Prevedel, R.; Ferrer Ortas, J.; Kerr, J. N.; Waters, J.; Breckwoldt, M. O.; Deneen, B.; Monje, M.; Soyka, S. J.; Venkataramani, V. Nature Reviews Neuroscience 2025, 1–17.

(22) Ferrer Ortas, J.; Mahou, P.; Escot, S.; Stringari, C.; David, N. B.; Bally-Cuif, L.; Dray, N.; Négrerie, M.; Supatto, W.; Beaurepaire, E. Light: Science & Applications 2023, 12, 29.

(23) Débarre, D.; Beaurepaire, E. Biophysical journal 2007, 92, 603–612.

(24) Bustamante-Marin, X. M.; Ostrowski, L. E. Cold Spring Harbor perspectives in biology 2017, 9, a028241.

(25) Abrami, M.; Biasin, A.; Tescione, F.; Tierno, D.; Dapas, B.; Carbone, A.; Grassi, G.; Conese, M.; Di Gioia, S.; Larobina, D., et al. International Journal of Molecular Sciences 2024, 25, 1933.

(26) Van Huizen, L. M.; Kuzmin, N. V.; Barbé, E.; van der Velde, S.; Te Velde, E. A.; Groot, M. L. Journal of biophotonics 2019, 12, e201800297.

(27) Murkar, R. S.; Wiese-Rischke, C.; Weigel, T.; Kopp, S.; Walles, H. Journal of Tissue Engineering 2025, 16, 20417314241299076.

(28) Becker, M. E.; Martin-Sancho, L.; Simons, L. M.; McRaven, M. D.; Chanda, S. K.; Hultquist, J. F.; Hope, T. J. Nature communications 2024, 15, 9480.

(29) Vishnu, K.; Subburaj, K.; Colombo, M. Computer Methods in Biomechanics and Biomedical Engineering 2025, 1–12.

(30) Sleigh, M. A.; Blake, J. R.; Liron, N. American Review of Respiratory Disease 1988, 137, 726–741.

(31) Bartlett, B. A.; Feng, Y.; Fromen, C. A.; Versypt, A. N. F. Computers & chemical engineering 2023, 179, 108458.

(32) Modaresi, M. A. Biomechanics and Modeling in Mechanobiology 2023, 22, 253–269.

(33) Hill, D. B.; Vasquez, P. A.; Mellnik, J.; McKinley, S. A.; Vose, A.; Mu, F.; Henderson, A. G.; Donaldson, S. H.; Alexis, N. E.; Boucher, R. C., et al. PloS one 2014, 9, e87681.

(34) Button, B.; Cai, L.-H.; Ehre, C.; Kesimer, M.; Hill, D. B.; Sheehan, J. K.; Boucher, R. C.; Rubinstein, M. Science 2012, 337, 937–941.

(35) Roth, D.; Şahin, A. T.; Ling, F.; Tepho, N.; Senger, C. N.; Quiroz, E. J.; Calvert, B. A.; van der Does, A. M.; Güney, T. G.; Glasl, S., et al. Nature communications 2025, 16, 2446.

(36) Ermund, A.; Meiss, L. N.; Rodriguez-Pineiro, A. M.; Bähr, A.; Nilsson, H. E.; Trillo-Muyo, S.; Ridley, C.; Thornton, D. J.; Wine, J. J.; Hebert, H., et al. Biochemical and biophysical research communications 2017, 492, 331–337.

(37) Fakih, D.; Rodriguez-Pineiro, A. M.; Trillo-Muyo, S.; Evans, C. M.; Ermund, A.; Hansson, G. C. American Journal of Physiology-Lung Cellular and Molecular Physiology 2020, 318, L1270–L1279.

(38) Donaldson, S. H.; Bennett, W. D.; Zeman, K. L.; Knowles, M. R.; Tarran, R.; Boucher, R. C. New England Journal of Medicine 2006, 354, 241–250.

(39) Reeves, E. P.; Molloy, K.; Pohl, K.; McElvaney, N. G. The Scientific World Journal 2012, 2012, 465230.

(40) Pangeni, R.; Meng, T.; Poudel, S.; Sharma, D.; Hutsell, H.; Ma, J.; Rubin, B. K.; Longest, W.; Hindle, M.; Xu, Q. International journal of pharmaceutics 2023, 634, 122661.

(41) Lai, S. K.; Wang, Y.-Y.; Hanes, J. Advanced drug delivery reviews 2009, 61, 158–171.

(42) Rogers, T. D.; Ostrowski, L. E.; Livraghi-Butrico, A.; Button, B.; Grubb, B. R. Scientific reports 2018, 8, 14744.

(43) Horton, N. G.; Wang, K.; Kobat, D.; Clark, C. G.; Wise, F. W.; Schaffer, C. B.; Xu, C. Nature photonics 2013, 7, 205–209.

(44) Li, B.; Li, J.; Zeb, J.; Yuan, Q.; Gan, W. Chemical Physics 2023, 572, 111955.

(45) Kim, D.; Mandl, G. A.; Balkota, M.; Vernaz, J.; Huang, S.; Constant, S.; Maechler, P.; DeWolf, C.; Capobianco, J. A.; Bonacina, L. Advanced Functional Materials 2024, 34, 2310078.

(46) Lin, H.; Li, H.; Cho, H.-J.; Bian, S.; Roh, H.-J.; Lee, M.-K.; Kim, J. S.; Chung, S.-J.; Shim, C.-K.; Kim, D.-D. Journal of pharmaceutical sciences 2007, 96, 341–350.

(47) Abraham, T.; Hirota, J. A.; Wadsworth, S.; Knight, D. A. Pulmonary pharmacology & therapeutics 2011, 24, 487–496.

(48) Grafström, R. C.; Nymark, P.; Hongisto, V.; Spjuth, O.; Ceder, R.; Willighagen, E.; Hardy, B.; Kaski, S.; Kohonen, P. Alternatives to Laboratory Animals 2015, 43, 325–332.

(49) Campargue, G.; La Volpe, L.; Giardina, G.; Gaulier, G.; Lucarini, F.; Gautschi, I.; Le Dantec, R.; Staedler, D.; Diviani, D.; Mugnier, Y., et al. Nano letters 2020, 20, 8725–8732.

(50) Ershov, D.; Phan, M.-S.; Pylvänäinen, J. W.; Rigaud, S. U.; Le Blanc, L.; Charles-Orszag, A.; Conway, J. R.; Laine, R. F.; Roy, N. H.; Bonazzi, D., et al. Nature methods 2022, 19, 829–832.

