## Supplementary Material for "Third Harmonic Generation Microscopy Reveals Structure and Mucus Dynamics in Human Airway Epithelium Models"

### Table of Contents

Video 1. Tracking of mucins and cellular debris in the mucus layer at different heights above the epithelial base: (a) 55 $\mu$m, (b) 65 $\mu$m, (c) 75 $\mu$m and (d) 85 $\mu$m.

Video 2. Time-lapse orthogonal view showing the THG signal from mucus and epithelium, corresponding to the region depicted in Fig. 3.

Supplementary Video 1. Maximum intensity projections of the mucus layer of epithelial tissue from CF-patient following apical application of PBS: (a) 0 $\mu$L, (b) 5 $\mu$L, and (c) 10 $\mu$L.

Supplementary Video 2. Maximum intensity projection and corresponding orthogonal views (along the dashed line) of epithelial tissue from CF patients following apical nebulization of 6% NaCl solution supplemented with Cy5 dye.

Supplementary Video 3. (a) Maximum intensity projection of MucilAir prior to the application of FBs; (b) 2 hours post-application.

Supplementary Video 4. (a) Maximum intensity projection of MucilAir prior to the application of FBs; (b) 20 hours post-application.

Supplementary Video 5. Maximum intensity projection and orthogonal view of lung epithelial tissue sequentially scanned under a 1 MHz OPA source (left) and a 80 MHz OPO source (right) at the same epithelial region over 21 minutes.

Supplementary Video 6. Brightfield video of the scanned region shown in Fig. S6 and its nearby area where the scanned region (white dashed line) and its adjacent area (red dashed line) exhibit no ciliary beating after 80 MHz OPA scanning.

Figure S1. THG image of the mucus layer

Figure S2. THG signal of fixed MucilAir after artificial removal of the overlying mucus layer under air exposure and after apical addition of PBS

Figure S3. THG image of the cilia layer and its orthogonal view in the absence of staining of epithelium or mucus

Figure S4. Experimental setup for aerosol deposition

Figure S5. THG and Cy5 fluorescence imaging of lung epithelial tissue during nebulization of hypertonic saline

Figure S6. Comparison of epithelial imaging using 1 MHz OPA and 80 MHz OPO excitation sources

Figure S7. Nonlinear microscope configuration used for harmonic generation microscopy

Table S1. Summary of donor characteristics and tissue properties of MucilAir and CF-MucilAir models used in this study

| Model | Donor | Pathology | Cell Type | Cilial Beating Frequency [Hz] | Tissue Integrity TEER $[\Omega \cdot cm^2]$ | Figure |
|---|---|---|---|---|---|---|
| MucilAir | A | No pathology reported | Bronchial | 7.1 ± 0.2 | 337 ± 06 | 1, S1 |
| | B | | Bronchial | 9.2 ± 0.3 | 356 ± 10 | 2 |
| | C | | Bronchial | 8.6 ± 0.2 | 332 ± 07 | 3, V1, V2 |
| | D | | Bronchial | 11.3 ± 0.1 | 261 ± 05 | S2 |
| | E | | Bronchial | 6.3 ± 0.2 | 1100 ± 10 | S3 |
| | F | | Bronchial | 8.1 ± 0.1 | 649 ± 12 | SV1, SV2 |
| | G | | Bronchial | 9.8 ± 0.3 | 451 ± 19 | SV3 |
| | H | | Bronchial | 8.7 ± 0.2 | 388 ± 15 | SV4 |
| | I | | Bronchial | 5.4 ± 0.1 | 471 ± 21 | SV5, SV6 |
| | J | | Bronchial | 6.9 ± 0.1 | 413 ± 09 | - |
| | K | | Bronchial | 8.4 ± 0.2 | 363 ± 12 | - |
| | L | | Bronchial | 7.8 ± 0.4 | 455 ± 25 | - |
| CF-MucilAir | M | Cystic Fibrosis | Bronchial | 9.4 ± 0.1 | 496 ± 09 | 4, SV1 |
| | N | | Bronchial | 6.0 ± 0.1 | 434 ± 07 | SV2 |

Table S1: Summary of donor characteristics and tissue properties of MucilAir and CF-MucilAir models used in this study.

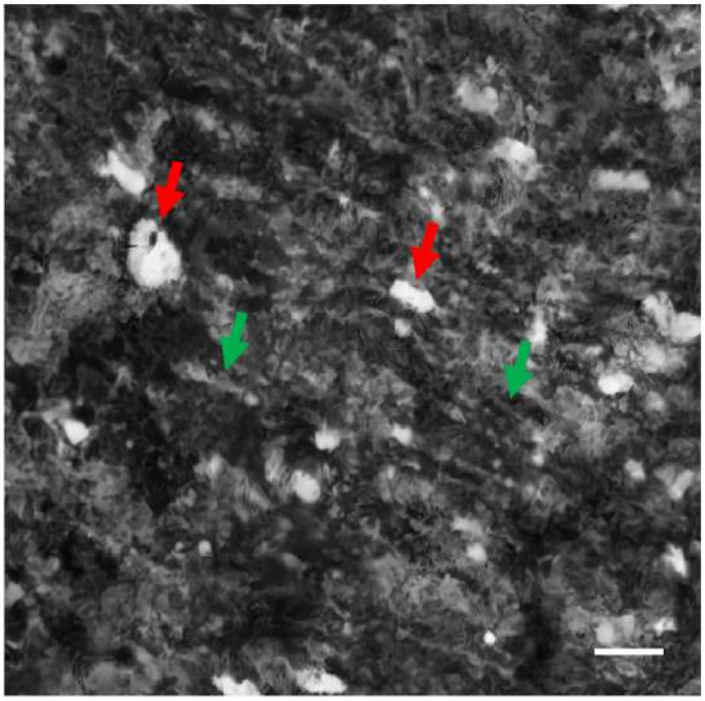

Figure S1: THG image of the mucus layer, showing filamentous (green arrow) and high contrast (red arrow) structures within the mucus. Scale bar: 20 $\mu$m

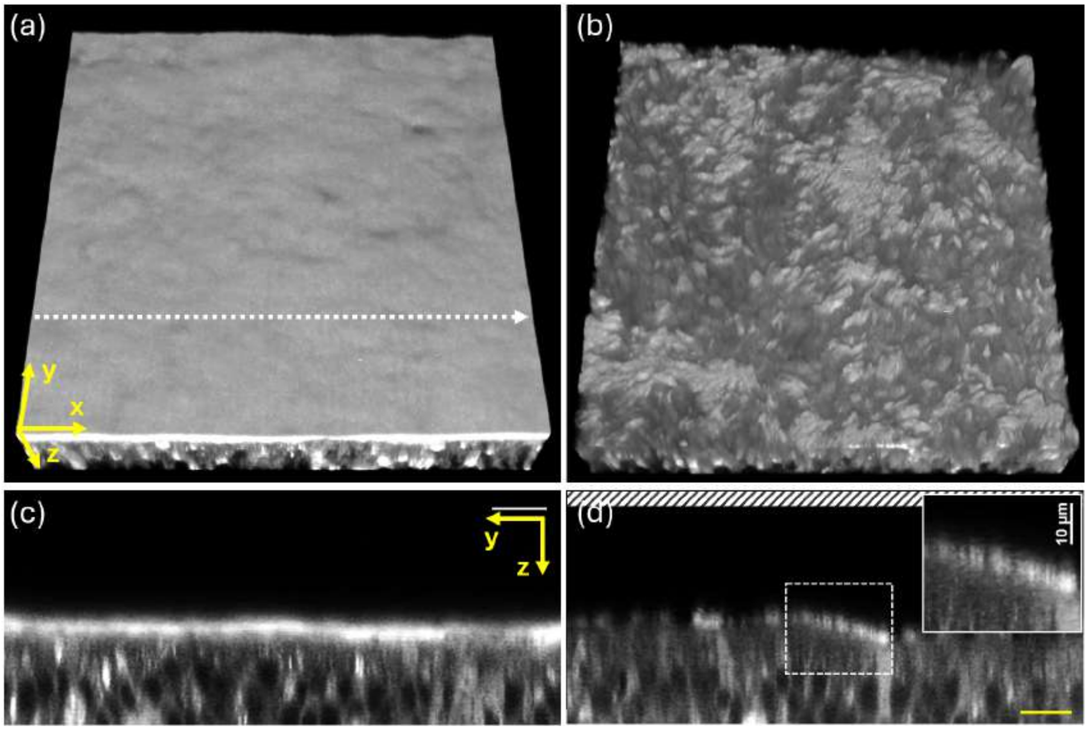


Figure S2: THG signal of fixed MucilAir after artificial removal of the overlying mucus layer, shown under two conditions: (a) when the epithelium is exposed to air, and (b) after apical addition of PBS, with (c–d) the corresponding orthogonal views along the dashed line. Inset in (d): magnified view of the boxed region. Scale bar: 20 $\mu$m; inset: 10 $\mu$m.

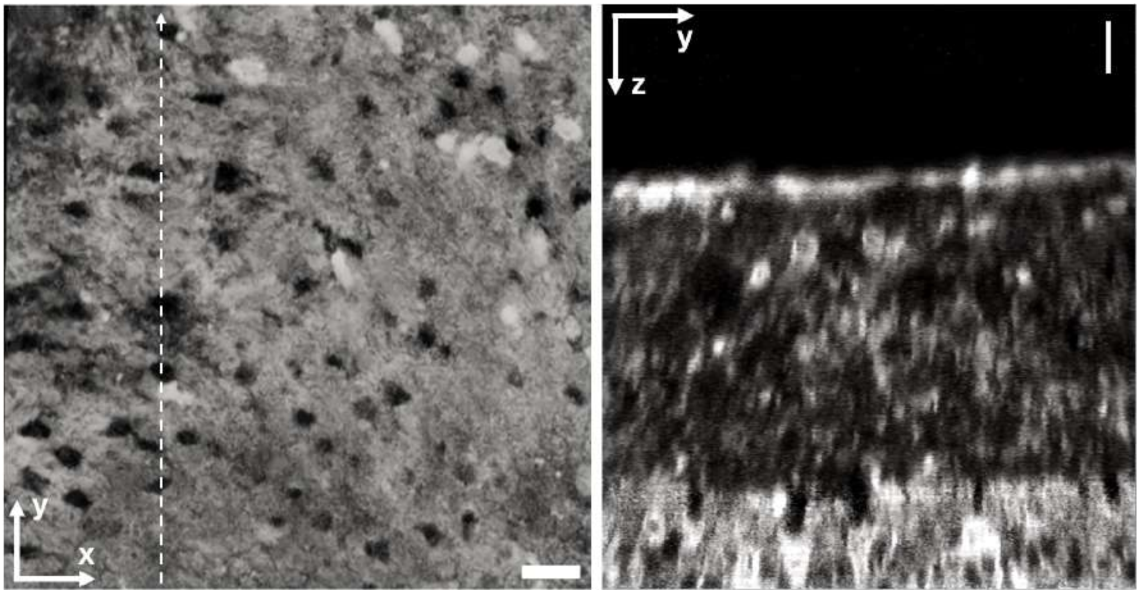


Figure S3: THG image of the cilia layer(left) and its orthogonal view along the dashed line (right), in the absence of staining procedure of epithelium or mucus. Scale bar: 20 $\mu$m

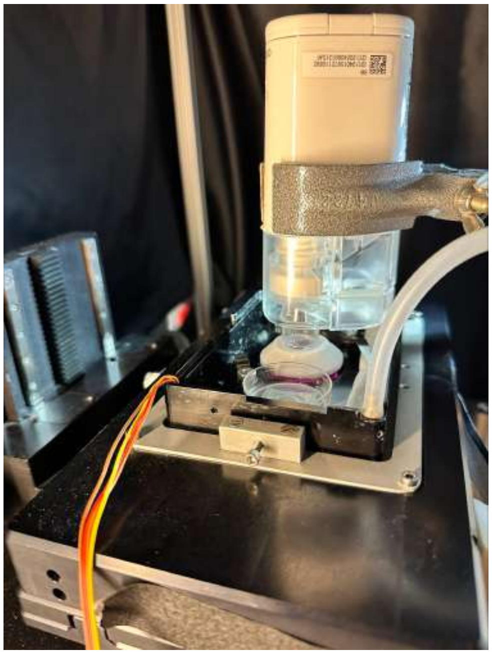

Figure S4: Set-up for aerosol deposition of 6 % NaCl solution with dissolved Cy5 dye (final concentration 1 $\mu$g/mL) using Omron NE-U100 on the apical side of MucilAir insert on the microscope stage.

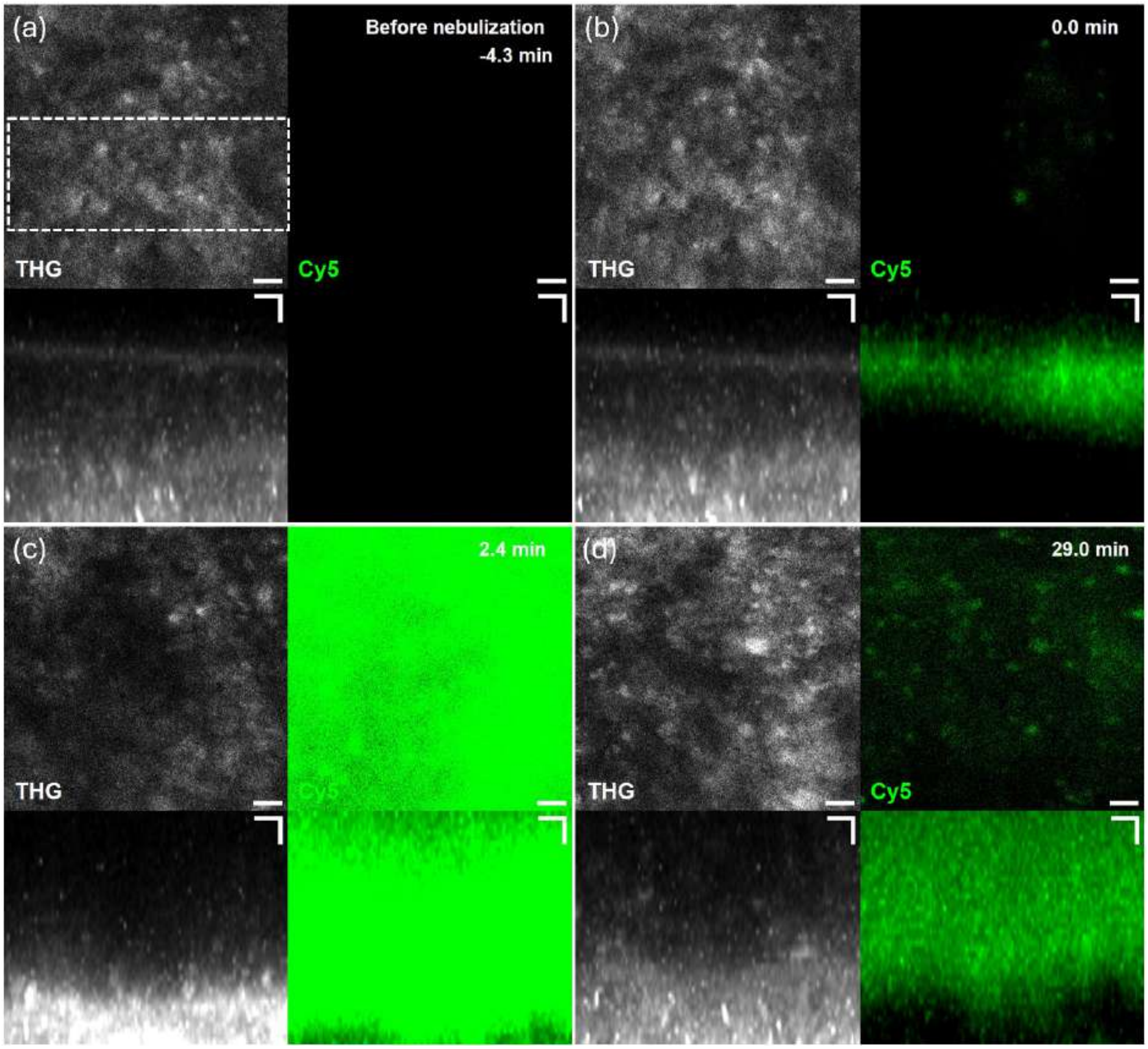


Figure S5: Maximum intensity projections of THG (grayscale) and Cy5 fluorescence (green) signals (top) and their corresponding maximum-intensity orthogonal projections (bottom) acquired from lung epithelial tissue derived from a CF patient during nebulization of hypertonic saline (6 % NaCl) supplemented with Cy5. THG provides label-free contrast of structural interfaces, while Cy5 reports the distribution of the fluorescent dye within the mucus layer. Maximum intensity projections depict the mucus layer only, whereas orthogonal views include both the epithelium and the mucus. Orthogonal projections were generated by maximum-intensity projection over the dashed region along the lateral dimension of the field of view. (a) Before nebulization. (b) initiation of nebulization, showing the initial appearance of fluorescence at the mucus surface. (c) Immediately after nebulization, with widespread Cy5 signal indicating penetration of the dye through the mucus layer. (d) 29 minutes after initiation of nebulization, showing redistribution and partial clearance of the dye. Note that the air–mucus interface is not visible in (c) because the orthogonal view is displayed over a fixed axial range, and the interface falls outside this range at later time points. Scale bar: 20 $\mu$m.

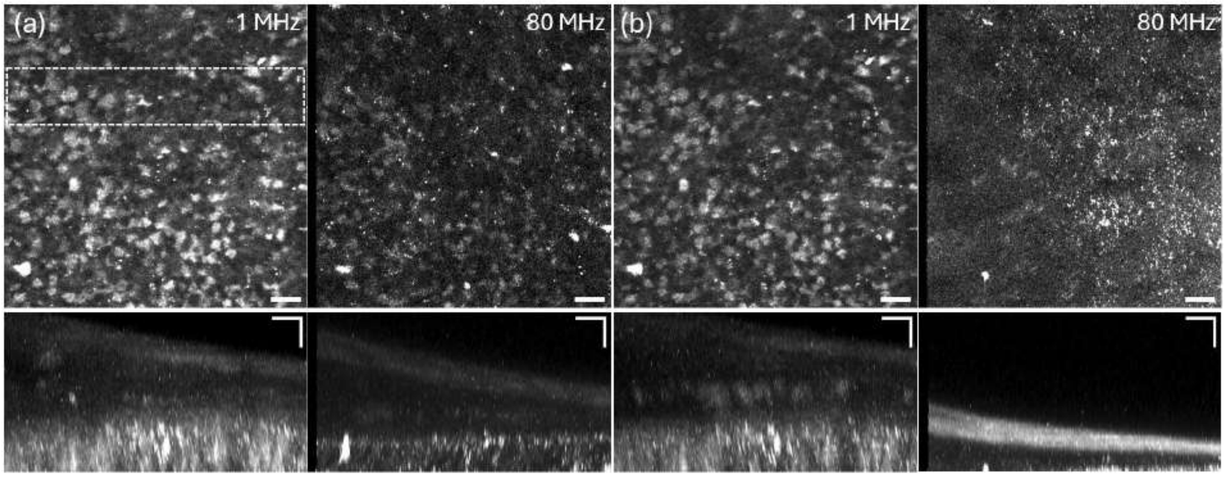


Figure S6: Maximum intensity projection (top) and orthogonal view (bottom) of lung epithelial tissue acquired in the same epithelial region using a 1 MHz OPA and an 80 MHz OPO at a frame rate of 1.55 Hz. The maximum intensity projections depict the epithelial layer only, whereas the orthogonal views include both the epithelium and the overlying mucus layer. Orthogonal projections were generated by maximum-intensity projection over the dashed region along the lateral dimension of the field of view. (a) Images acquired immediately after the first scan and (b) images acquired 21 minutes after sequential scanning under identical conditions. Note that the first 5 $\mu$m of each OPO line scan were not exposed to the excitation beam due to the response delay of the external shutter (see Fig. S7). Scale bar: 20 $\mu$m.

**Qualitative comparison of low- and high-repetition-rate excitation regimes**

As shown in Fig. S6, the same epithelial region was sequentially imaged using the 1 MHz OPA and the 80 MHz OPO. The experiment with the 1 MHz OPA was performed first to avoid potential tissue damage, followed by a 30-minute resting period before acquisition with the 80 MHz OPO to reduce potential thermal effects. While the 1 MHz OPA laser did not cause detectable damage to the epithelial tissue or the overlying mucus layer, the 80 MHz OPO source induced immediate and localized tissue damage in the same region, accompanied by drying of the mucus layer. This damage is also shown in Supplementary Video 6, recorded under brightfield microscopy, where the scanned region (dashed line) and its nearby area (red dashed line) exhibit no ciliary beating following exposure.

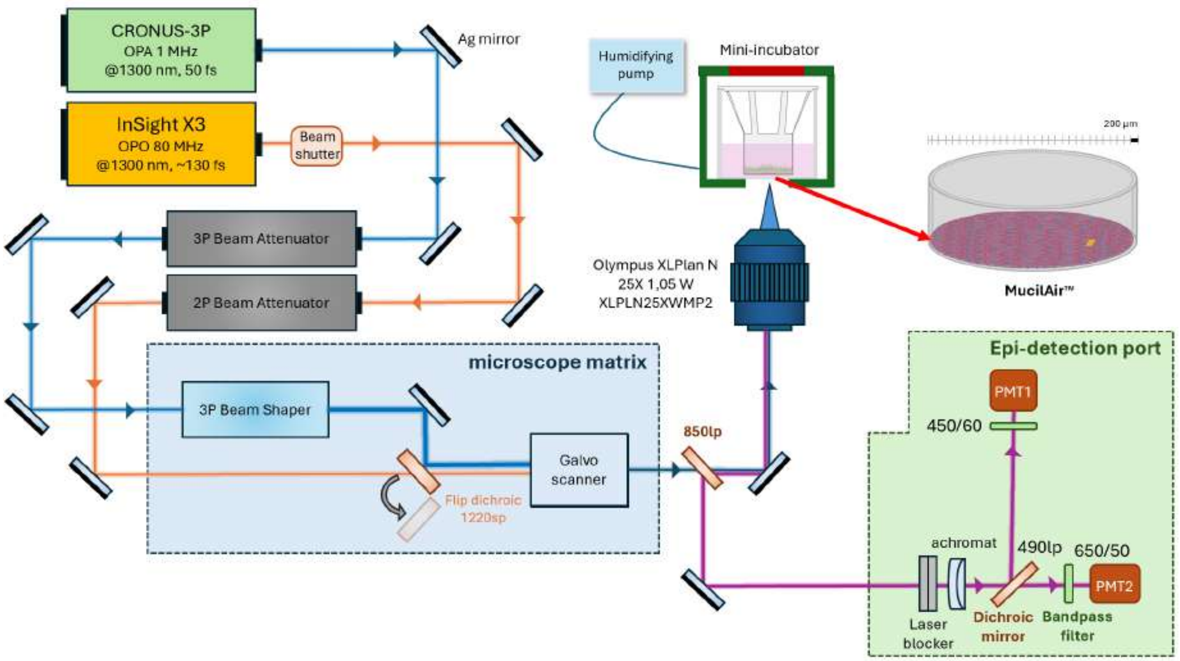


Figure S7: Nonlinear microscope configuration. Two femtosecond excitation sources, the CRONUS-3P (OPA, 1 MHz, 1250–1800 nm) and Insight X3 (OPO, 80 MHz, 680–1300 nm), were attenuated and, for the OPA line, beam-shaped before before being relayed to a galvo–galvo scanner. A 1220 nm short pass dichroic mirror directed the excitation beam within the microscope matrix (blue box) and was flipped out for InSight X3 operation at 1300 nm. The excitation beam and emitted signals were separated with an 850 nm long pass dichroic. Signals were detected with two photomultiplier tubes (PMT1: Hamamatsu H12056-40; PMT2: Hamamatsu H12056-20). Spectral separation of THG and fluorescence was achieved using a 490 long pass dichroic mirror together with 450/60 nm and 650/50 nm bandpass filters. A fast shutter was inserted in the InSight X3 beam path to block the beam during flyback and beam parking. The CRONUS-3P was externally triggered to blank the beam during the same interval. The dimensions of the scanning field of view and the inset are indicated in the top right in yellow (200 × 200 $\mu m^2$).
