## Supplementary figures and images for "Third Harmonic Generation Microscopy Reveals Structure and Mucus Dynamics in Human Airway Epithelium Models"

### Fig. S1

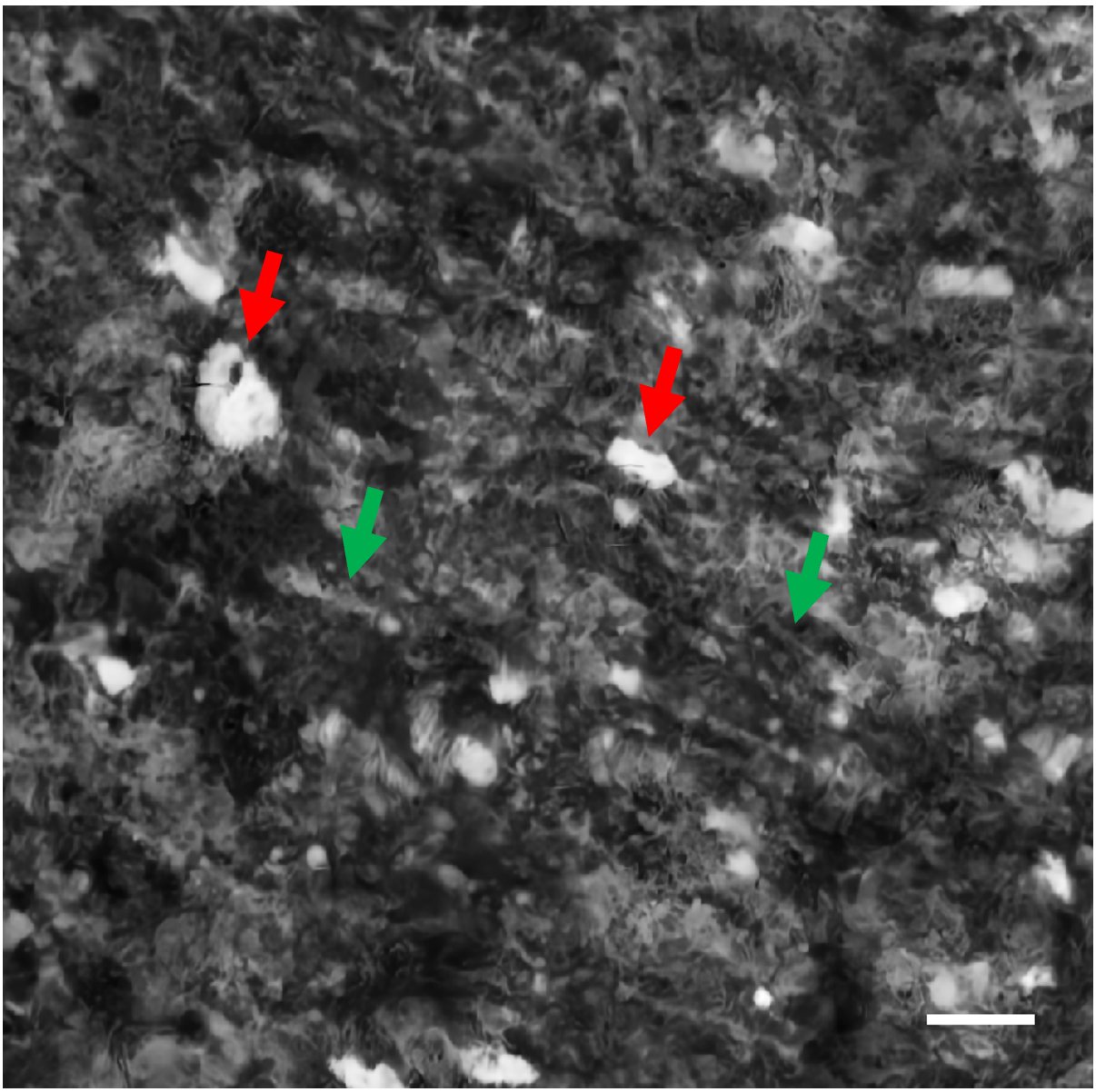

### Fig. S2

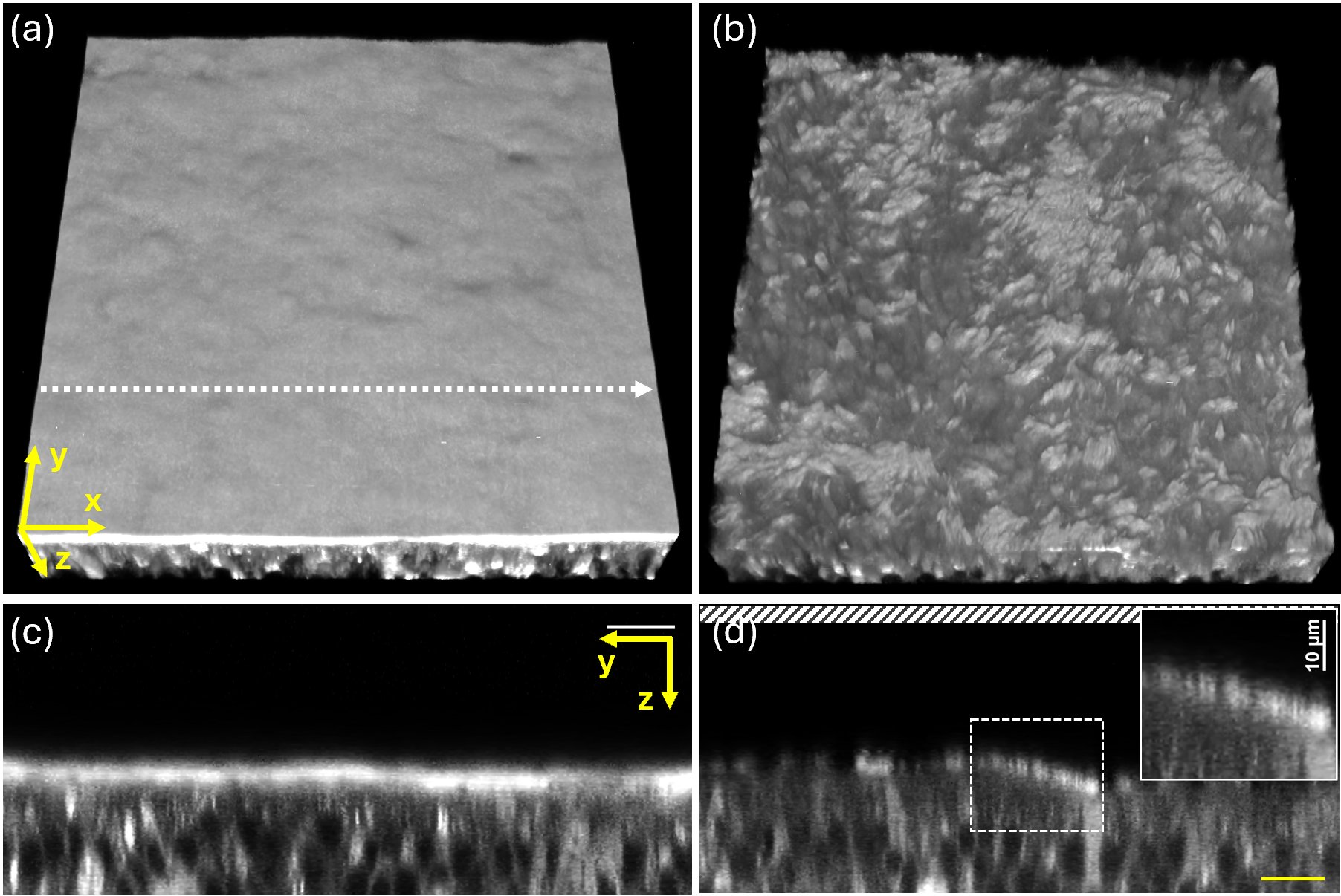

### Fig. S3

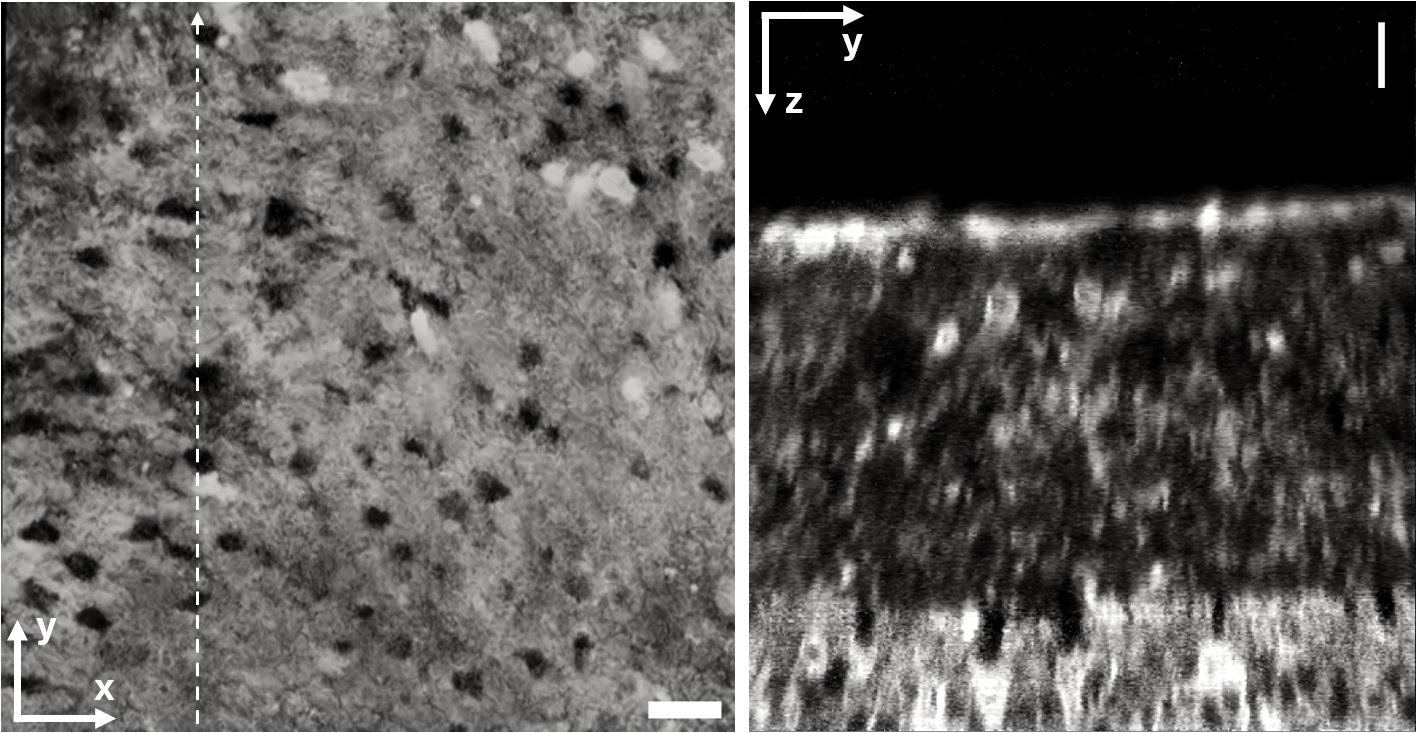

### Fig. S4

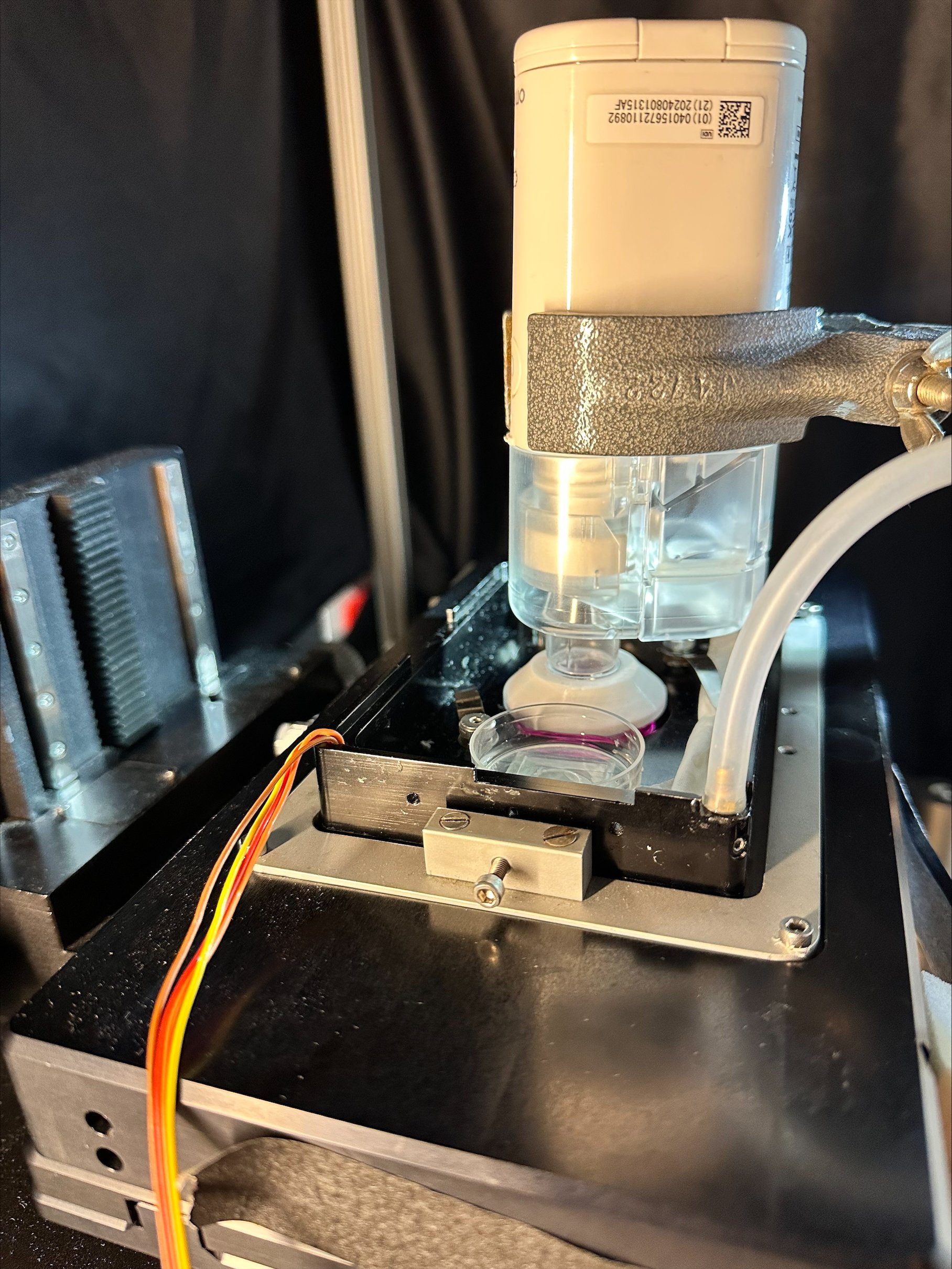

### Fig. S5

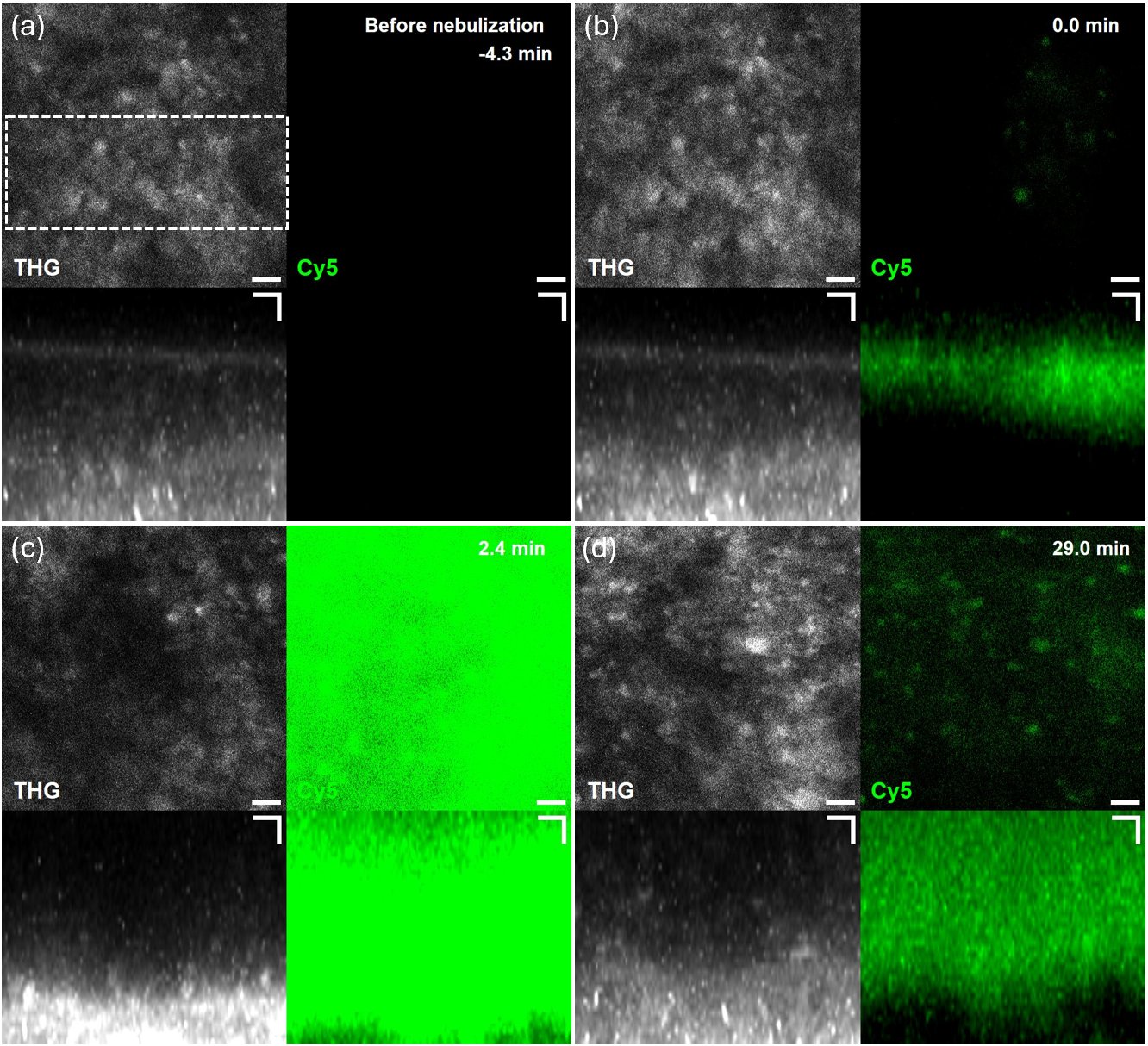

### Fig. S6

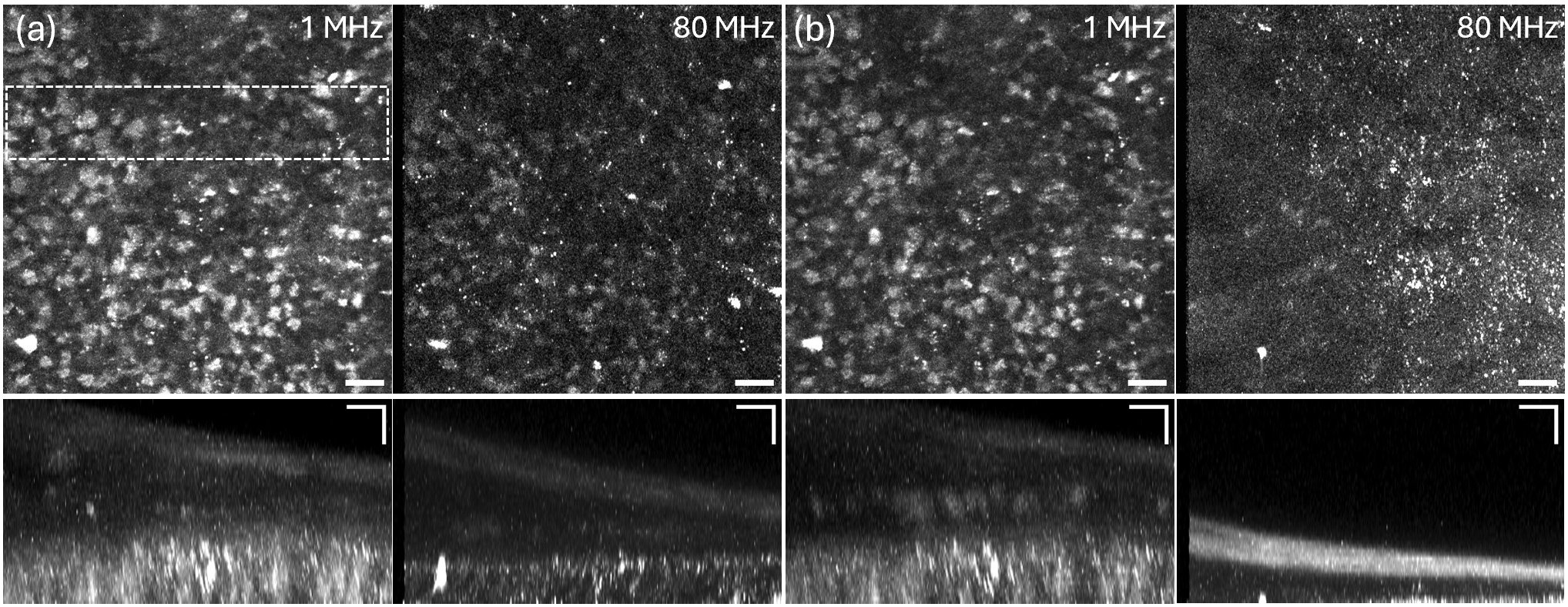

### Fig. S7

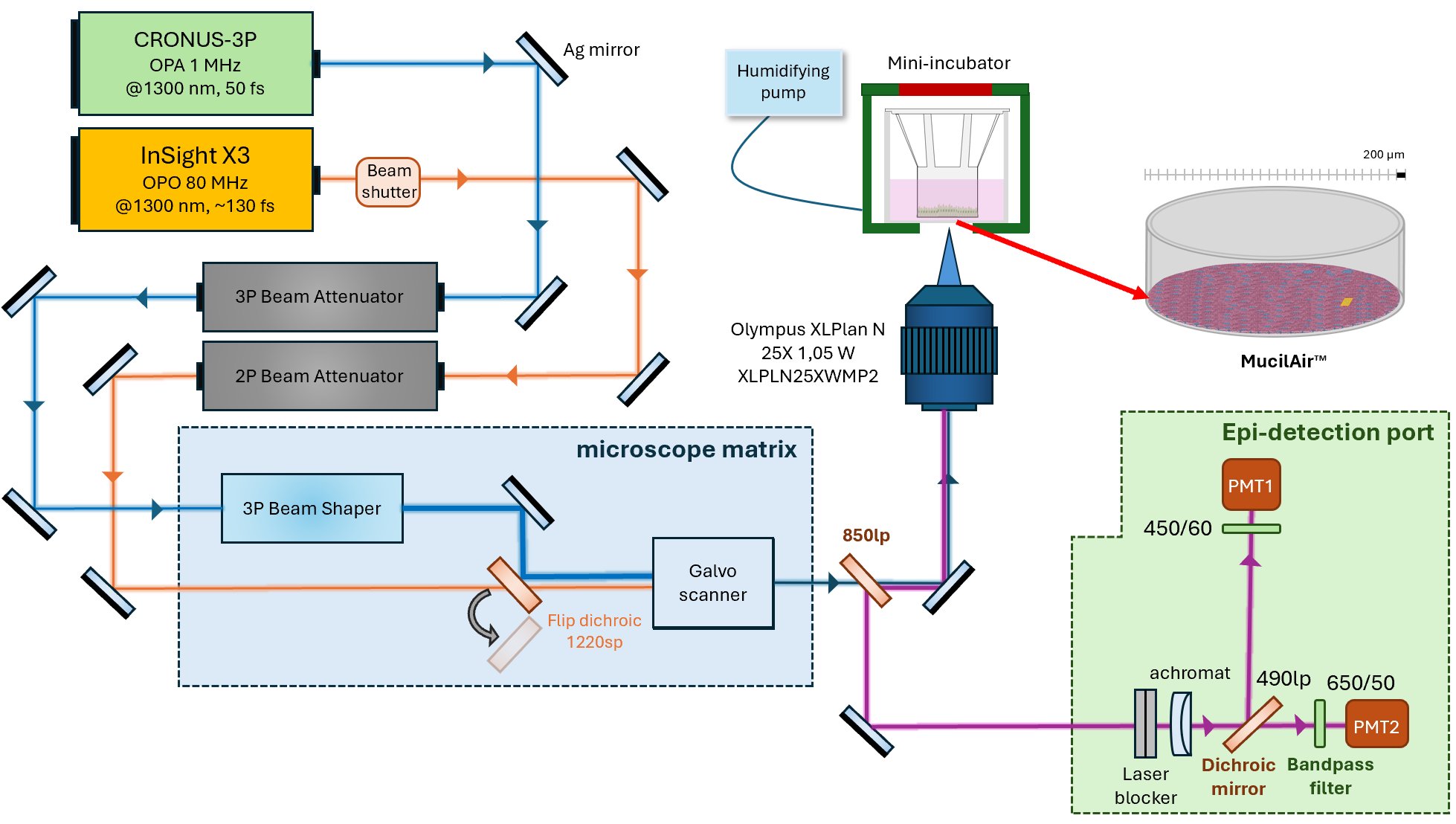
